# Incentive salience attribution, “sensation-seeking” and “novelty-seeking” are independent traits in a large sample of male and female heterogeneous stock rats

**DOI:** 10.1101/421065

**Authors:** Alesa R. Hughson, Aidan P. Horvath, Katie Holl, Abraham A. Palmer, Leah C. Solberg Woods, Terry E. Robinson, Shelly B. Flagel

## Abstract

There are a number of traits that are thought to increase susceptibility to addiction, and some of these are modeled in preclinical studies. For example, “sensation-seeking” is predictive of the initial propensity to take drugs; whereas “novelty-seeking” predicts compulsive drug-seeking behavior. In addition, the propensity to attribute incentive salience to reward cues can predict the propensity to approach drug cues, and reinstatement or relapse, even after relatively brief periods of drug exposure. The question addressed here is the extent to which these three ‘vulnerability factors’ are related; that is, predictive of one another. Some relationships have been reported in small samples, but here a large sample of 1,598 outbred male and female heterogeneous stock rats were screened for Pavlovian conditioned approach behavior (to obtain an index of incentive salience attribution; ‘sign-tracking’), and subsequently tested for sensation-seeking and novelty-seeking. Despite the large N there were no significant correlations between these traits, in either males or females. There were, however, novel relationships between multiple measures of incentive salience attribution and, based on these findings, we generated a new metric that captures “incentive value”. Furthermore, there were sex differences on measures of incentive salience attribution and sensation-seeking behavior that were not previously apparent.

## Introduction

Stimuli (‘cues’) in the environment can guide adaptive behavior, bringing an individual into close proximity to valuable resources (e.g. food, water, mates) or leading one away from danger. However, for some individuals, cues associated with reward can gain excessive control over behavior and lead to maladaptive outcomes^1^. In human drug addicts, cues that have been previously associated with the drug-taking experience can themselves acquire the ability to motivate drug-seeking behavior and can instigate relapse, even when there is a expressed desire to discontinue use ^2,3^. The ability of reward-associated cues to motivate behavior occurs largely through Pavlovian learning processes^4-6^. When a previously neutral stimulus is repeatedly paired with presentation of a reward, it acquires predictive properties and becomes a conditioned stimulus (CS), and in some cases also acquires incentive motivational properties, and thus the ability to act as an incentive stimulus^7-9^.

There is, however, considerable individual variation in the extent to which animals attribute incentive motivational value (“incentive salience”) to reward-associated cues^9-12^. For some rats, known as “goal-trackers” (GTs), the reward cue serves only as a predictor (a CS) and evokes a conditioned response (CR) directed toward the location of reward delivery^10^. For others, termed “sign-trackers” (STs), the CS is both predictive and attractive, and evokes a CR directed towards the CS ^11,13^. Importantly, relative to GTs, STs are also more motivated to self-administer cocaine^14^, more likely to approach drug cues, and exhibit enhanced cue- and drug-induced reinstatement of cocaine-seeking behavior after relatively limited drug exposure and a brief period of abstinence^14,15^. These data support the notion that differences in the way individuals learn about Pavlovian cue-reward associations are applicable to the study of drug abuse and addiction.

Other traits that have been associated with addiction vulnerability include the propensity to engage in “sensation-seeking” and “novelty-seeking” behaviors^16-21^. These are, undoubtedly, multidimensional traits, but with considerable conceptual and empirical overlap, at least in humans^22^. In rats, “sensation-seeking” is assessed via locomotor response to an inescapable novel environment^23^; whereas “novelty-seeking” is indicated by preference for a novel environment when given a choice (i.e. novelty-induced conditioned place preference)^24^. The sensation-seeking trait has been shown to be a good predictor of the initial propensity to take drugs in rodents^25^; whereas novelty-seeking better predicts the propensity for compulsive drug use^26^. While these two traits appear to represent distinct facets of addiction vulnerability, the relationship between them is not well understood. There are some reports of a negative correlation between sensation-seeking and novelty-seeking behavior^27,28^ and others indicating no relationship^29^. These inconsistencies could be due to a number of factors including, differences in the testing paradigms, order of testing and sample size.

We, and others, have sought to determine whether the propensity to attribute incentive salience to reward cues represents another divergent addiction-related trait, or if it is related to either sensation-seeking or novelty-seeking behavior. In outbred Sprague-Dawley rats, there is no apparent correlation between locomotor response to an inescapable novel environment and the propensity to attribute incentive salience to reward cues^9,27,30^. Yet, in rats that are selectively bred based on sensation-seeking behavior, these traits are highly correlated^31^, likely due to the selective-breeding paradigm and resultant combination of traits inherent to these phenotypes^32^. It should be noted, however, that these selectively-bred rats do not differ in novelty-seeking behavior^32^. The relationship between the propensity to attribute incentive salience to reward cues and novelty-seeking behavior, has, to our knowledge, been examined in just one study in outbred rats. Beckmann et al. ^27^ reported a positive relationship between these two traits, but with a relatively small sample size. Thus, further investigation is needed to elucidate the relationship between these addiction-related traits.

In the current study, we used a large sample of heterogeneous stock (HS) rats to further explore the relationship between: 1) the propensity to attribute incentive salience to reward cues, 2) sensation-seeking behavior, and 3) novelty-seeking behavior. HS rats were created by combining eight inbred strains together and subsequently maintaining the colony in a way that minimizes inbreeding^33^. The generation of HS rats from a single breeding colony helps avoid any spurious correlations between traits that may arise as a result of population structure^34^. These rats, therefore, serve as a unique and invaluable model to investigate the relationship between multiple addiction-related endophenotypes. Male and female HS rats were assessed for the propensity to attribute incentive salience to reward cues using a Pavlovian conditioned approach paradigm followed by a conditioned reinforcement test. Rats were subsequently tested for locomotor response to novelty (i.e. sensation-seeking) and novelty-preference (i.e. novelty-seeking). The relationship between these three traits was then examined in a population of 1,598 rats, with sex as an independent variable.

## Results

To illustrate group comparisons, several of the following datasets are displayed using notched box plots. A box plot is illustrated in Figure 1, given that some readers will not be familiar with them. The boundaries of the box represent the interquartile range, the horizontal line between the notches indicates the median, and the width of the notch represents a 95% confidence interval around the median. Thus, if the notches do not overlap this indicates a group difference with 95% confidence. The vertical line within the box represents the standard error of the mean (the mean would be at the mid-point of this line).

**Figure 1.**
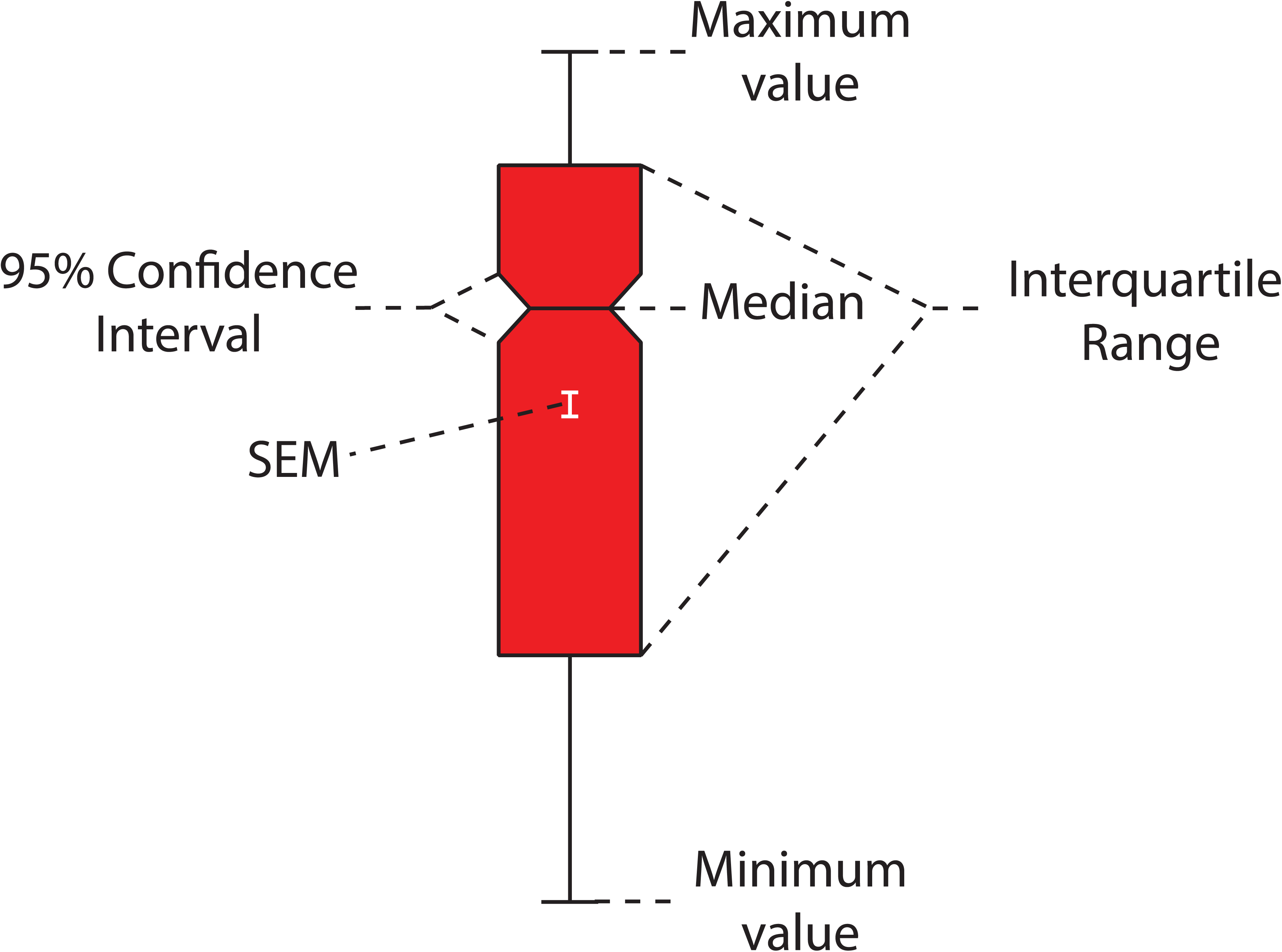
Notched box plot summary. An example of a notched box plot with labels for each informational aspect as described in the text. SEM, standard error of the mean.

### Pavlovian conditioned approach index score in male and female HS rats

Figure 2 illustrates Pavlovian conditioned approach (PCA) behavior for female (n=799) and male (n=799) HS rats as measured by their PCA Index score. The PCA Index score was calculated from a number of metrics of approach to the food cup or the lever, as previously described^35^. Briefly, a score of -1 is indicative of behavior directed exclusively towards the food cup (i.e. an extreme goal-tracker), and +1 is indicative of behavior directed exclusively towards the lever (i.e. an extreme sign-tracker). Over the course of training the average PCA Index score increased, from that reflective of mainly goal-tracking (presumably because rats had been pretrained to retrieve food from the food cup) to more reflective of sign-tracking (effect of session, F_4, 1596_=269.6, p<0.001; Figure 2a). In addition, there was a significant effect of sex (F_1, 1599.357_=47.580, p<0.001) and a significant interaction between sex and session (F_4, 1596_=25.00, p<0.001; Figure 2a). The PCA Index was greater for males relative to females on the first session of PCA training (p<0.005), but on all subsequent sessions was greater in females, and this effect became more pronounced as training progressed (Session 2-5, p<0.001). In agreement, there was a significant sex difference in the terminal (average of sessions 4-5) PCA Index score (F_1, 1597_=59.87, p<0.001; Cohen’s d=0.42), as shown in Figure 2b, where there is a perceptible gap between the 95% confidence intervals for females vs. males. Furthermore, the frequency histogram in Figure 2c further illustrates the bias towards sign-tracking behavior in females compared to males. Taken together, these results suggest that females, as a group, have a greater propensity to attribute incentive salience to a food cue, at least as assessed by sign-tracking behavior.

**Figure 2.**
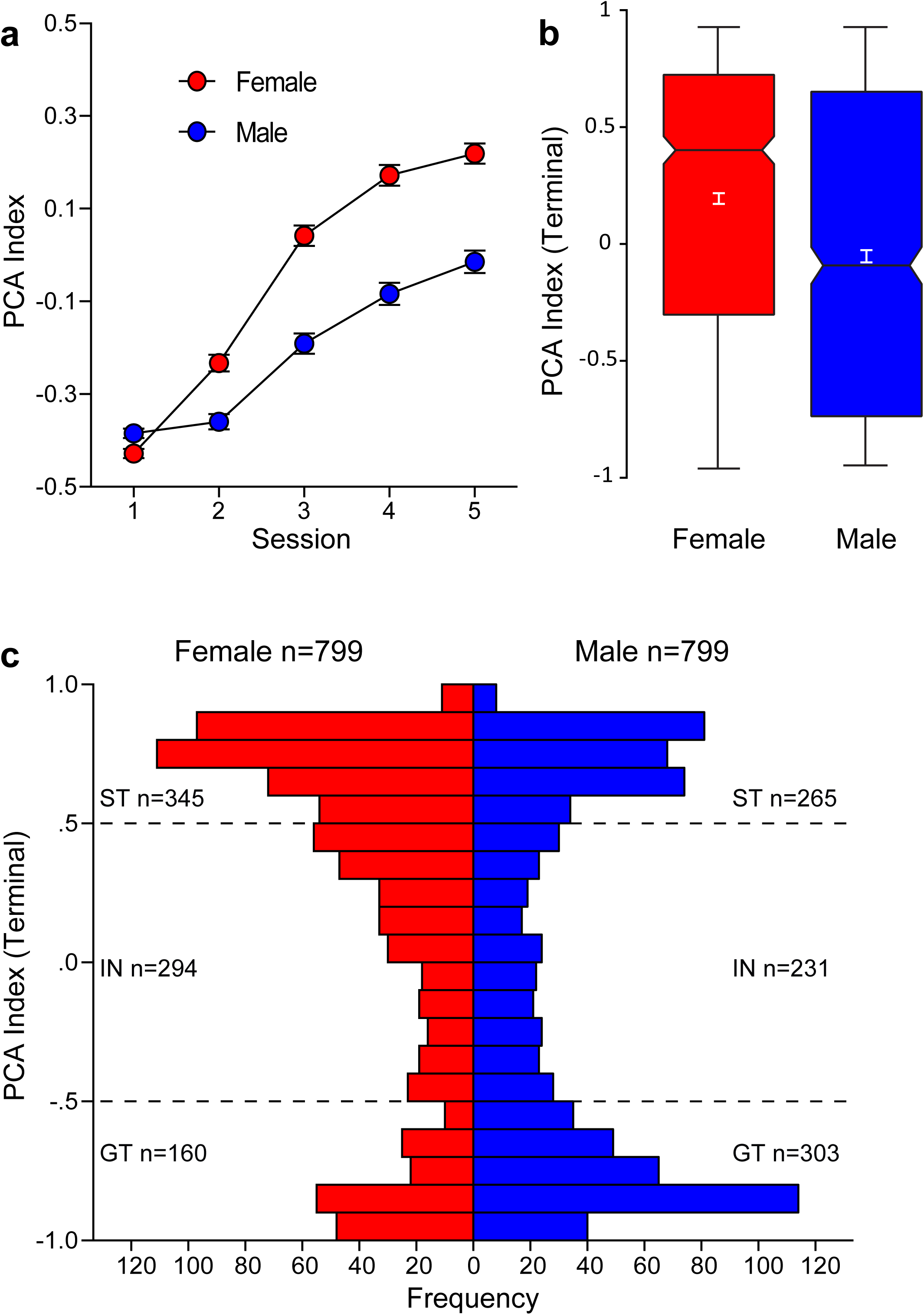
Pavlovian conditioned approach Index score in male and female HS rats. A Pavlovian conditioned approach (PCA) Index score was calculated for each rat as described in the text. A score of - 1 indicates exclusive food cup-oriented behavior and a score of +1 indicates exclusive lever-oriented behavior. (a) PCA Index mean ± SEM for each of 5 training sessions for female (n=799) and male (n=799) rats. (b) Notched box plot of terminal PCA Index, calculated as the average of the PCA Index from sessions 4 and 5, for male and female rats. (c) Histogram of the distribution of terminal PCA Index for females and males. Horizontal hashed lines indicate the threshold of PCA Index used to define phenotype groups (GT, -5 – 0; intermediate responders -0.5 - +0.5; STs, +0.5 – 1).

### Sign-tracking and goal-tracking behavior

Rats were characterized as STs, intermediate responders (INs) or GTs based on their terminal PCA Index score, as described previously^35^, and differences between phenotypes were assessed across the 5 Pavlovian training sessions for various measures of lever-directed/sign-tracking behavior and food cup-directed/goal-tracking behavior (Figure 3). Main effects and interactions from the statistical analyses are reported in Table 1. For each metric, there was a significant effect of session, indicating a change in behavior over the course of training, as rats acquired their respective CRs. In addition, there was a significant effect of phenotype and sex for all metrics. As expected, STs learned to direct their behavior towards the lever-conditioned stimulus (CS) to a greater extent than INs and GTs, displaying a higher probability (Figure 3a), increased vigor (Figure 3c), and decreased latency (Figure 3e) to deflect the lever-CS. These differences were apparent from the second session of training onward. INs had a greater tendency towards sign-tracking behavior relative to GTs. Conversely, GTs learned to approach the food cup during the CS period. As shown in Figure 3b, d, and f, GTs approached the food cup with higher probability, increased vigor, and decreased latency compared to the other two phenotypes. INs were more likely than STs to enter the food cup during lever-CS presentation. There was a significant three-way interaction between session, sex, and phenotype for all of these metrics except Food Cup Entry Probability, as indicated in Table 1. Bonferroni-corrected posthoc comparisons between phenotypes for each session are listed in Table 2; and comparisons between sexes within each phenotype are listed in Table 3. On measures of sign-tracking behavior, the most robust sex differences appear to be in the IN phenotype, such that female INs exhibit greater sign-tracking behavior across sessions relative to male INs. In contrast, male and female GTs differ on measures of goal-tracking behavior, with female GTs showing more robust behavior directed toward the food cup across sessions. Although differences between sexes were less apparent in STs, the enhanced responding in both female INs and female GTs compared to their male counterparts suggests that these differences could be due to greater baseline activity levels in females. This notion is further supported by enhanced responding at the food cup during the inter-trial interval in females compared to males, for each session of training (averaged across sessions: female, 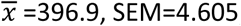; males, 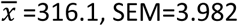). However, given the robust sex differences displayed in the terminal PCA Index score (Figure 2b), it is unlikely that increased activity levels alone can account for the enhanced sign-tracking behavior observed in females.

**Figure 3.**
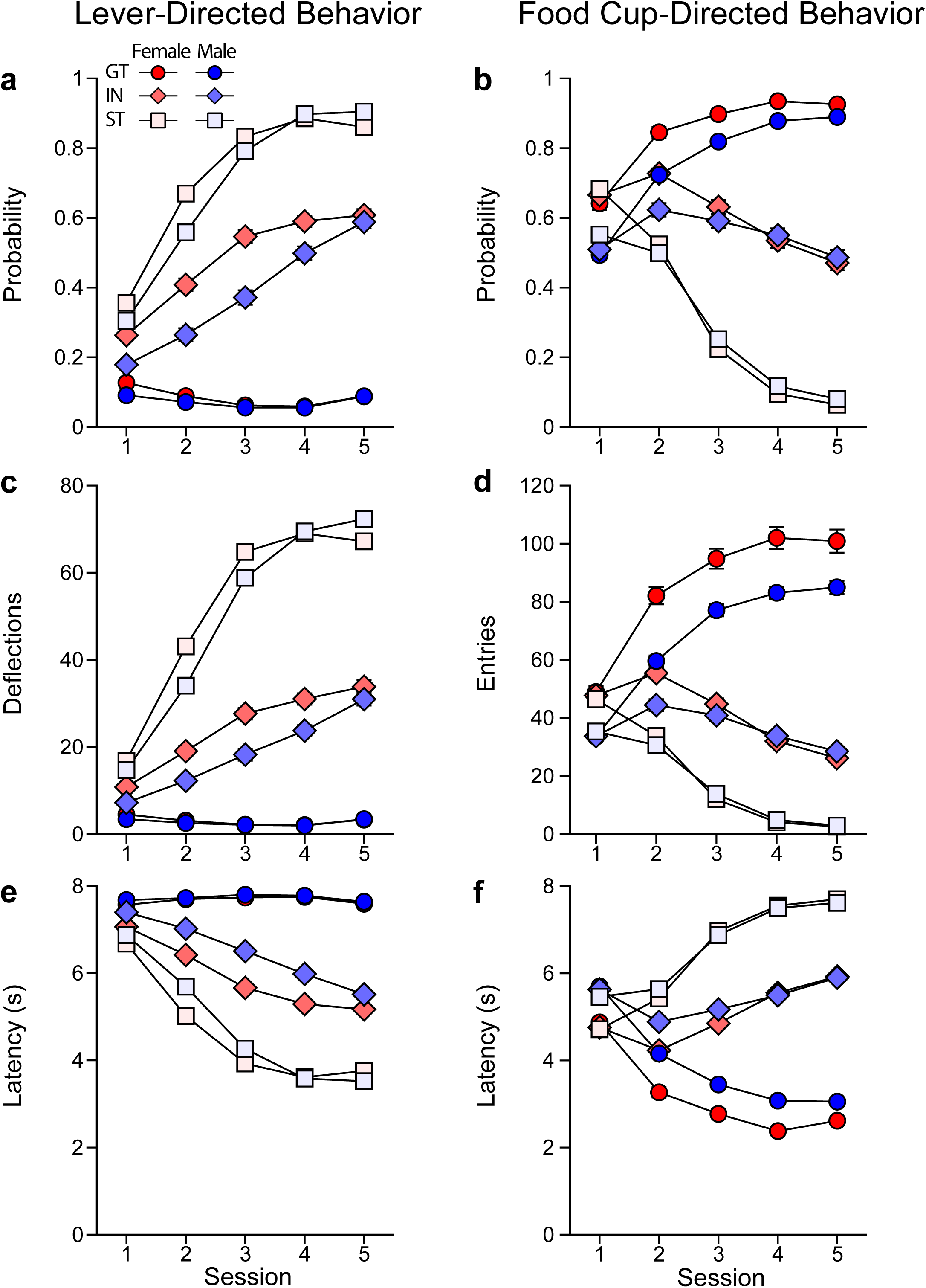
Sign-tracking and goal-tracking behavior. Behavior displayed for the 5 sessions of PCA training for each phenotype of each sex (Female GT (n=160), IN (n=294), ST (n=345); Male GT (n=303), IN (n=231), ST (n=265)) as mean ± SEM of the following metrics: (a) probability to deflect the lever; (b) probability to enter the food cup; (c) number of lever deflections; (d) number of food cup entries; (e) latency to deflect the lever, and (f) latency to enter the food cup.

**Table 1.** Results from the linear mixed model analyses for sign-tracking (top) and goal-tracking (bottom) behaviors. The effects of sex, phenotype and session and interactions were analyzed. Abbreviations: DF, degrees of freedom, SS, sum of squares

| Sign-tracking Behaviors | Lever Deflection Probability |  |  | Lever Deflections |  |  | Lever Latency |  |  |
| --- | --- | --- | --- | --- | --- | --- | --- | --- | --- |
|  | DF,SS | F-Value | P-Value | DF,SS | F-Value | P-Value | DF,SS | F-Value | P-Value |
| Effect of Sex | 1, 1602 | 32.26 | <0.001 | 1, 1591 | 11.90 | <0.005 | 1, 1592 | 29.70 | <0.001 |
| Effect of Phenotype | 2, 1602 | 1778 | <0.001 | 2, 1591 | 1184 | <0.001 | 2, 1592 | 1234 | <0.001 |
| Sex*Phenotype | 2, 1602 | 10.44 | <0.001 | 2, 1591 | 3.915 | <0.005 | 2, 1592 | 9.311 | <0.001 |
| Effect of Session | 4, 2204 | 586.5 | <0.001 | 4, 1591 | 572.9 | <0.001 | 4, 1592 | 709.0 | <0.001 |
| Session*Sex | 4, 2204 | 12.78 | <0.001 | 4, 1591 | 7.455 | <0.001 | 4, 1592 | 8.54 | <0.001 |
| Session*Phenotype | 8, 2204 | 224.5 | <0.001 | 8, 1591 | 275.8 | <0.001 | 8, 1592 | 259.2 | <0.001 |
| Session*Sex*Phenotype | 8, 2204 | 4.906 | <0.001 | 8, 1591 | 3.623 | <0.001 | 8, 1592 | 5.95 | <0.001 |

| Goal-tracking Behaviors | Food Cup Entry Probability |  |  | Food Cup Entries |  |  | Food Cup Latency |  |  |
| --- | --- | --- | --- | --- | --- | --- | --- | --- | --- |
|  | DF,SS | F-Value | P-Value | DF,SS | F-Value | P-Value | DF,SS | F-Value | P-Value |
| Effect of Sex | 1, 1592 | 34.18 | <0.001 | 1, 1592 | 59.90 | <0.001 | 1, 1592 | 58.71 | <0.001 |
| Effect of Phenotype | 2, 1592 | 994.0 | <0.001 | 2, 1592 | 941.2 | <0.001 | 2, 1592 | 1085 | <0.001 |
| Sex*Phenotype | 2, 1592 | 5.002 | <0.01 | 2, 1592 | 18.91 | <0.001 | 2, 1592 | 9.351 | <0.001 |
| Effect of Session | 4, 1592 | 132.4 | <0.001 | 4, 1592. | 59.11 | <0.001 | 4, 1592 | 111.6 | <0.001 |
| Session*Sex | 4, 1592 | 25.29 | <0.001 | 4, 1592. | 8.302 | <0.001 | 4, 1592 | 19.16 | <0.001 |
| Session*Phenotype | 8, 1592 | 327.5 | <0.001 | 8, 1592. | 260.8 | <0.001 | 8, 1592 | 359.7 | <0.001 |
| Session*Sex*Phenotype | 8, 1592 | 1.906 | 0.055 | 8, 1592. | 4.593 | <0.001 | 8, 1592 | 4.061 | <0.001 |

**Table 2.** Posthoc comparisons between phenotypes for each session of Pavlovian conditioning for sign-tracking (top) and goal-tracking (bottom) behaviors. Abbreviations: GT, goal-tracker; IN, intermediate responder; ST, sign-tracker

| Sign-tracking Behaviors |  | Lever Deflection Probability |  |  |  |  |
| --- | --- | --- | --- | --- | --- | --- |
| Phenotype Comparison |  | Session 1 | Session 2 | Session 3 | Session 4 | Session 5 |
| GT vs. IN |  | p<0.001* | p<0.001* | p<0.001* | p<0.001* | p<0.001* |
| GT vs. ST |  | p<0.001* | p<0.001* | p<0.001* | p<0.001* | p<0.001* |
| IN vs. ST |  | p<0.001* | p<0.001* | p<0.001* | p<0.001* | p<0.001* |
|  |  | Lever Deflections |  |  |  |  |
|  |  | Session 1 | Session 2 | Session 3 | Session 4 | Session 5 |
| GT vs. IN |  | p<0.001* | p<0.001* | p<0.001* | p<0.001* | p<0.001* |
| GT vs. ST |  | p<0.001* | p<0.001* | p<0.001* | p<0.001* | p<0.001* |
| IN vs. ST |  | p<0.001* | p<0.001* | p<0.001* | p<0.001* | p<0.001* |
|  |  | Lever Latency |  |  |  |  |
|  |  | Session 1 | Session 2 | Session 3 | Session 4 | Session 5 |
| GT vs. IN |  | p<0.001* | p<0.001* | p<0.001* | p<0.001* | p<0.001* |
| GT vs. ST |  | p<0.001* | p<0.001* | p<0.001* | p<0.001* | p<0.001* |
| IN vs. ST |  | p<0.001* | p<0.001* | p<0.001* | p<0.001* | p<0.001* |
| Goal-tracking Behaviors |  | Food Cup Entry Probability |  |  |  |  |
| Phenotype Comparison |  | Session 1 | Session 2 | Session 3 | Session 4 | Session 5 |
| GT vs. IN |  | p=0.546 | p<0.001* | p<0.001* | p<0.001* | p<0.001* |
| GT vs. ST |  | p<0.005* | p<0.001* | p<0.001* | p<0.001* | p<0.001* |
| IN vs. ST |  | p=0.084 | p<0.001* | p<0.001* | p<0.001* | p<0.001* |
|  |  | Food Cup Entries |  |  |  |  |
|  |  | Session 1 | Session 2 | Session 3 | Session 4 | Session 5 |
| GT vs. IN |  | p=1 | p<0.001* | p<0.001* | p<0.001* | p<0.001* |
| GT vs. ST |  | p=1 | p<0.001* | p<0.001* | p<0.001* | p<0.001* |
| IN vs. ST |  | p=1 | p<0.001* | p<0.001* | p<0.001* | p<0.001* |
|  |  | Food Cup latency |  |  |  |  |
|  |  | Session 1 | Session 2 | Session 3 | Session 4 | Session 5 |
| GT vs. IN |  | p=0.801 | p<0.001* | p<0.001* | p<0.001* | p<0.001* |
| GT vs. ST |  | p<0.05* | p<0.001* | p<0.001* | p<0.001* | p<0.001* |
| IN vs. ST |  | p=0.520 | p<0.001* | p<0.001* | p<0.001* | p<0.001* |

\* indicates significant difference between phenotypes following Bonferroni correction

**Table 3.** Posthoc comparisons between sexes for each phenotype on each session of Pavlovian conditioning for sign-tracking (top) and goal-tracking (bottom) behaviors. Abbreviations: M, male; F, female

| Sign-tracking Behaviors<br>Sex Comparison<br>(M vs F within each phenotype) |  | Lever Deflection Probability |  |  |  |  |
| --- | --- | --- | --- | --- | --- | --- |
|  |  | Session 1 | Session 2 | Session 3 | Session 4 | Session 5 |
|  | GT | p=0.077 | p=0.512 | p=0.789 | p=0.867 | p=0.982 |
|  | IN | p<0.001* | p<0.001* | p<0.001* | p<0.001* | p=0.274 |
|  | ST | p<0.005* | p<0.001* | p<0.05* | p=0.455 | p<0.05* |
|  |  | Lever Deflections |  |  |  |  |
|  |  | Session 1 | Session 2 | Session 3 | Session 4 | Session 5 |
|  | GT | p=0.378 | p=0.800 | p=0.979 | p=0.952 | p=0.966 |
|  | IN | p<0.005* | p<0.001* | p<0.001* | p<0.001* | p=0.179 |
|  | ST | p<0.05* | p<0.001* | p<0.005* | p=0.730 | p<0.01* |
|  |  | Lever Latency |  |  |  |  |
|  |  | Session 1 | Session 2 | Session 3 | Session 4 | Session 5 |
|  | GT | p=0.213 | p=0.856 | p=0.629 | p=0.816 | p=0.724 |
|  | IN | p<0.001* | p<0.001* | p<0.001* | p<0.001* | p<0.005* |
|  | ST | p<0.01* | p<0.001* | p<0.005* | p=0.769 | p<0.05* |
| Goal-tracking Behaviors<br>Sex Comparison<br>(M vs F within each phenotype) |  | Food Cup Entry Probability |  |  |  |  |
|  |  | Session 1 | Session 2 | Session 3 | Session 4 | Session 5 |
|  | GT | p<0.001* | p<0.001* | p<0.01* | p<0.05* | p=0.080 |
|  | IN | p<0.001* | p<0.001* | p=0.08 | p=0.416 | p=0.392 |
|  | ST | p<0.001* | p=0.254 | p=0.190 | p=0.208 | p=0.347 |
|  |  | Food Cup Entries |  |  |  |  |
|  |  | Session 1 | Session 2 | Session 3 | Session 4 | Session 5 |
|  | GT | p<0.001* | p<0.001* | p<0.001* | p<0.001* | p<0.001* |
|  | IN | p<0.001* | p<0.001* | p=0.140 | p=0.467 | p=0.335 |
|  | ST | p<0.001* | p=0.225 | p=0.437 | p=0.718 | p=0.868 |
|  |  | Food Cup latency |  |  |  |  |
|  |  | Session 1 | Session 2 | Session 3 | Session 4 | Session 5 |
|  | GT | p<0.001* | p<0.001* | p<0.001* | p<0.001* | p<0.001* |
|  | IN | p<0.001* | p<0.001* | p<0.05* | p=0.573 | p=0.778 |
|  | ST | p<0.001* | p=0.111 | p=0.474 | p=0.576 | p=0.362 |

\* indicates significant difference between sexes following Bonferroni correction

### Conditioned reinforcement

The conditioned reinforcement test revealed that the lever had reinforcing properties for all animals, as there was a greater number of responses into the active port (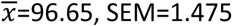) relative to the inactive port (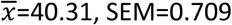) when phenotype and sex were collapsed (effect of port: F_1, 3180_=1184, p<0.001, Cohen’s d = 1.22). This effect was also apparent when the sexes were analyzed separately (females, effect of port, F_1, 1589_=930.2, p<0.001, Cohen’s d = 1.53; males, effect of port, F_1,1590_=423.3, p<0.001, Cohen’s d = 1.03; Figure 4a). There was also a significant port x sex interaction (F_1, 3180_=107.5, p<0.001). Although post-hoc comparisons revealed that, relative to males, females made significantly more nose pokes into both the active (p<0.001) and inactive (p<0.001) ports, the interaction appears to stem from a greater effect of sex in the active port (Cohen’s d = 0.79) compared to the inactive port (Cohen’s d = 0.41). Nonetheless, to account for significant sex differences in responding in the inactive port, responses in the inactive port were subtracted from those in the active port (A-I), and differences between sexes and phenotypes were assessed using this metric (Figure 4b). In agreement with the data above, females had significantly higher A-I scores compared to males (effect of sex: F_1, 1589_=152.6, p<0.001, Cohen’s d =0.62). In addition, there was a significant main effect of phenotype for both females (F_2, 793_=14.91, p<0.001) and males (F_2, 792_=18.63, p<0.001), such that, for both sexes, STs exhibited a greater A-I score relative to GTs (female: p<0.001; male: p<0.001). Additionally, neither female (p=0.025) nor male (p=0.069) STs differed in A-I score from INs, and INs had a higher score than GTs for both females (p<0.01) and males (p<0.005). In summary, these data indicate that the lever acts as a more effective conditioned reinforcer for STs compared to INs, and for INs compared to GTs, in both sexes, which is consistent with prior reports^9,36^.

**Figure 4.**
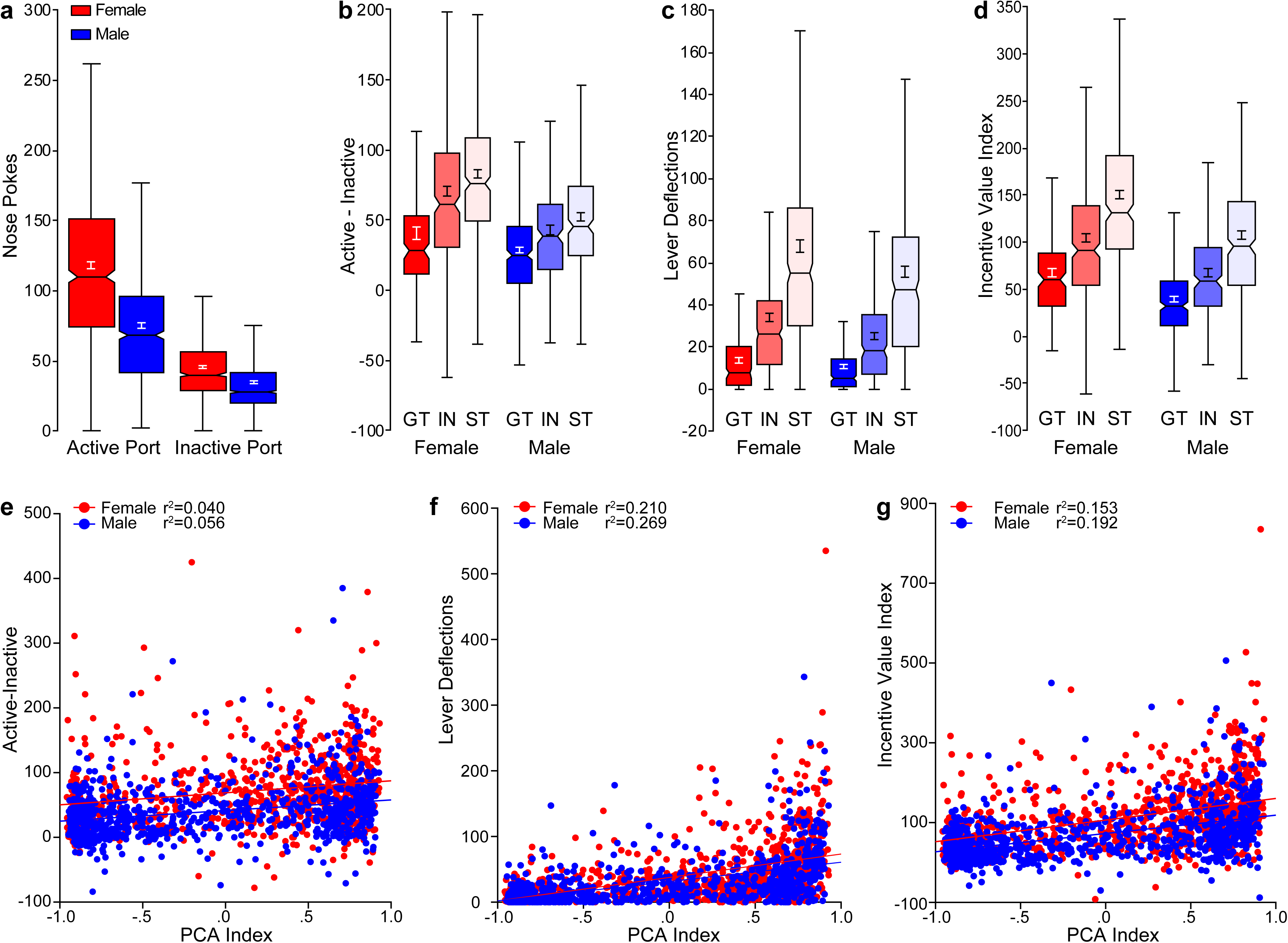
Conditioned reinforcement. (a) Notched box plot of the number of nose pokes into the active and inactive ports for both female (n=795) and male (n=795-796) rats. (b) Notched box plot of nose pokes into the active port minus nose pokes into the inactive port for females (GT (n=160), IN (n=293), ST (n=341)) and males (GT (n=302), IN (n=227), ST (n=264)). (c) Lever deflections during the conditioned reinforcement test for females (GT (n=160), IN (n=293), ST (n=341)) and males (GT (n=302), IN (n=228), ST (n=264)). (d) Notched box plot of Incentive Value Index for females (GT (n=160), IN (n=293), ST (n=341)) and males (GT (n=302), IN (n=227), ST (n=264)). (e) Plot of each individual rat’s data for Active minus Inactive (A-I) value as a function of terminal PCA Index with linear regression line. (f) Plot of each individual rat’s data for lever deflections as a function of terminal PCA Index with linear regression line. (g) Plot of each individual rat’s Incentive Value Index as a function of terminal PCA Index with linear regression line.

Following a nose poke into the active port, the lever was extended for 2 seconds. During these 2 seconds, it is not uncommon for the rats to manipulate the lever, especially if they are sign-trackers^9^. Figure 4c shows the number of lever deflections during the 2-second period it was available for each phenotype and sex. Consistent with the findings above, females made more lever deflections compared to males (effect of sex: F_1, 1590_=46.65, p<0.001); however, the size of this effect was relatively small (Cohen’s d = 0.34). There was a significant main effect of phenotype for both sexes (females, F_2, 793_=102.4, p<0.001; males, F_2, 793_=141.4, p<0.001). As expected, STs deflected the lever upon its presentation more than GTs or IN rats, and this was true for both sexes (female and male: ST, GT p<0.001; ST, IN p<0.001). In addition, relative to GTs, IN rats responded more on the lever for both sexes (female and male: IN, GT p<0.001). These data indicate that each phenotype engaged with the lever to a significantly different degree, with STs engaging most avidly, GTs engaging least avidly, and INs showing an intermediate level of interest.

The group differences in lever-directed behavior are important because behavior directed at the lever would compete with the ability to respond into the nose port and thereby result in underestimating the conditioned reinforcing properties of the lever, especially in STs. Thus, in order to account for both nose-port responding and lever deflections, we calculated a new metric, the “Incentive Value Index” ((responses in active port – responses in inactive port) + lever deflections)). For this outcome measure, there was a significant effect of sex (F_1, 1586_=94.25, p<0.001, Cohen’s d=0.60) and phenotype (F_2, 1586_=142.4, p<0.001; Figure 4d). Similar to the lever deflection results, females had a higher Incentive Value Index compared to males, STs had a higher score compared to both GTs (p<0.001) and INs (p<0.001), and INs had a higher score compared to GTs (p<0.001).

The ability of PCA behavior to predict the conditioned reinforcing properties of the lever was assessed using linear regression between the terminal PCA Index score and the A-I responses (Figure 4e), lever deflections (Figure 4f) and Incentive Value Index (Figure 4g). For all analyses, there was a main effect of sex on the dependent variable (A-I: F_1, 1587_=111.6, p<0.001; lever deflections: F_1, 1587_=10.55, p<0.005; Incentive Value Index: F_1,1586_=85.47, p<0.001). Only for lever deflections, however, was there a significant interaction between sex and PCA Index (F_1, 1587_=4.381, p<0.05). These data indicate that the sexes differed in their relationship between PCA score and lever-oriented behavior. All outcome measures of the conditioned reinforcement test were significantly (p<0.001) and positively correlated with PCA Index, but the size of the effect as assessed by the r^2^ value suggests that the relationship between A-I score and PCA Index is relatively weak, accounting for less than 10% of the variance (Both sexes r^2^=0.066; female r^2^=0.040; male r^2^=0.056). Lever deflections and Incentive Value Index were more strongly correlated with the PCA Index, accounting for ∼20-25% of the variance (lever deflections: both sexes r^2^=0.246; female r^2^=0.210; male r^2^=0.269; Incentive Value Index: both sexes r^2^=0.193; female r^2^=0.153; male r^2^=0.192). Taken together, these data suggest that interaction with the CS is a critical component of the conditioned reinforcement test and one that should be incorporated when assessing the incentive motivational value of reward cues.

### Sensation-seeking behavior: Locomotor response to novelty

Figure 5a shows the locomotor response to a novel environment for each sex and phenotype. There was a significant main effect of sex (F_1, 1529_=11.19, p<0.005), such that females travelled greater distances than males, and a significant effect of phenotype (F_2, 1529_=6.016, p<0.005); STs travelled further than GTs (p<0.005). There was not a significant difference in distance travelled between INs and either STs (p=0.058) or GTs (p=0.110). Figure 5b shows the total distance travelled following placement into a novel environment plotted as a function of terminal PCA Index score for each individual rat. For the regression, there was a significant main effect of sex (F_1, 1529_=10.38, p<0.005), but no interaction between sex and PCA Index (F_1, 1529_=0.648, p=0.421). Although the correlation between these two metrics was statistically significant (p<0.001), the size of the effect is too small to constitute any meaningful relationship (both sexes: r^2^=0.013; female: r^2^=0.006; male r^2^=0.013). Thus, it appears that an individual’s tendency to attribute incentive salience to a food cue is not related to “sensation-seeking” behavior, and this is true for both females and males.

**Figure 5.**
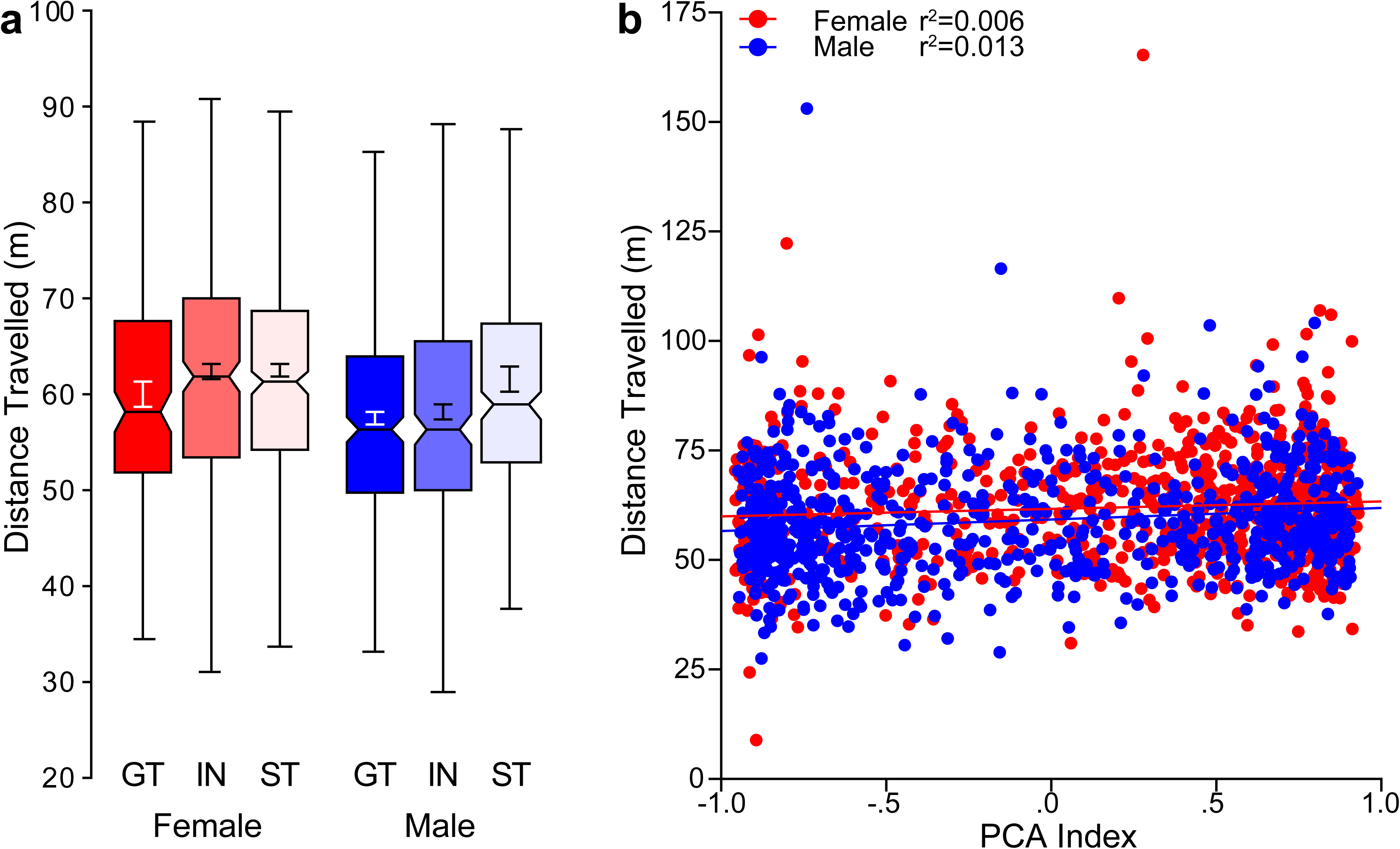
Locomotor response to novelty. Novelty-induced locomotor activity assessed on the first habituation day of the novelty-seeking paradigm. (a) Notched box plot of total distance (meters) travelled for females (GT (n=152), IN (n=280), ST (n=332)) and males (GT (n=292), IN (n=222), ST (n=252)). (b) Plot of each individual rat’s data for total distance travelled during habituation as a function of terminal PCA Index with linear regression line.

When the relationship between PCA Index score and sensation-seeking behavior is examined in a subset of the population stratified according to response to the novel environment, a slightly different pattern of results emerges. That is, when rats are divided based on a median spilt of the distance travelled, a significant regression was found in those that exhibited low levels of activity (“low-responders”; p<0.001), but not those that exhibited high levels of activity (“high-responders”; p=0.528). However, in both cases, the effect size was too small to constitute a meaningful relationship (high-responders both sexes: r^2^=0.0005; female: r^2^=0.002; male r^2^=0.003; low-responders both sexes: r^2^=0.034; female: r^2^=0.028; male r^2^=0.036).

### Novelty-seeking behavior: Novelty place preference

All rats spent more time in the novel zone of the test chamber relative to the familiar zone (F_1, 3041_=6.609, p<0.05). There was no effect of side bias with respect to how the chambers were configured in the testing room (chamber side, (F_1,139_=0.388, P=0.534). Moreover, the degree of zone preference was consistent between the sexes (sex x zone interaction, (F_1, 3041_=0.990, p=0.320); Figure 6b) and there were no significant differences between sexes (F_1, 1517_=1.645, p=0.200) or phenotypes (F_2, 1517_=0.089, p=0.914) for the percent of time spent in the novel zone during the test session (Figure 6c). To assess whether PCA behavior predicted novelty-seeking behavior, we plotted our metric of interest (% time spent in the novel zone) as a function of terminal PCA Index score for each individual rat with sex as a covariate (Figure 6d). In this case, there was not a significant main effect of sex (F_1, 1517_=1.557, p=0.212), nor was there a significant interaction between sex and PCA Index (F_1, 1517_=0.857, p=0.355), indicating that the relationship between traits was similar for each sex. Contrary to prior reports^27^, we did not find a significant correlation between these two traits (both sexes r^2^=0.0006, p=0.356; female r^2^=0.00007, p=0.819; male r^2^=0.001, p=0.288). Thus, novelty place preference appears to be unrelated to an individual’s tendency to attribute incentive salience to a food cue, and this is true for both males and females.

**Figure 6.**
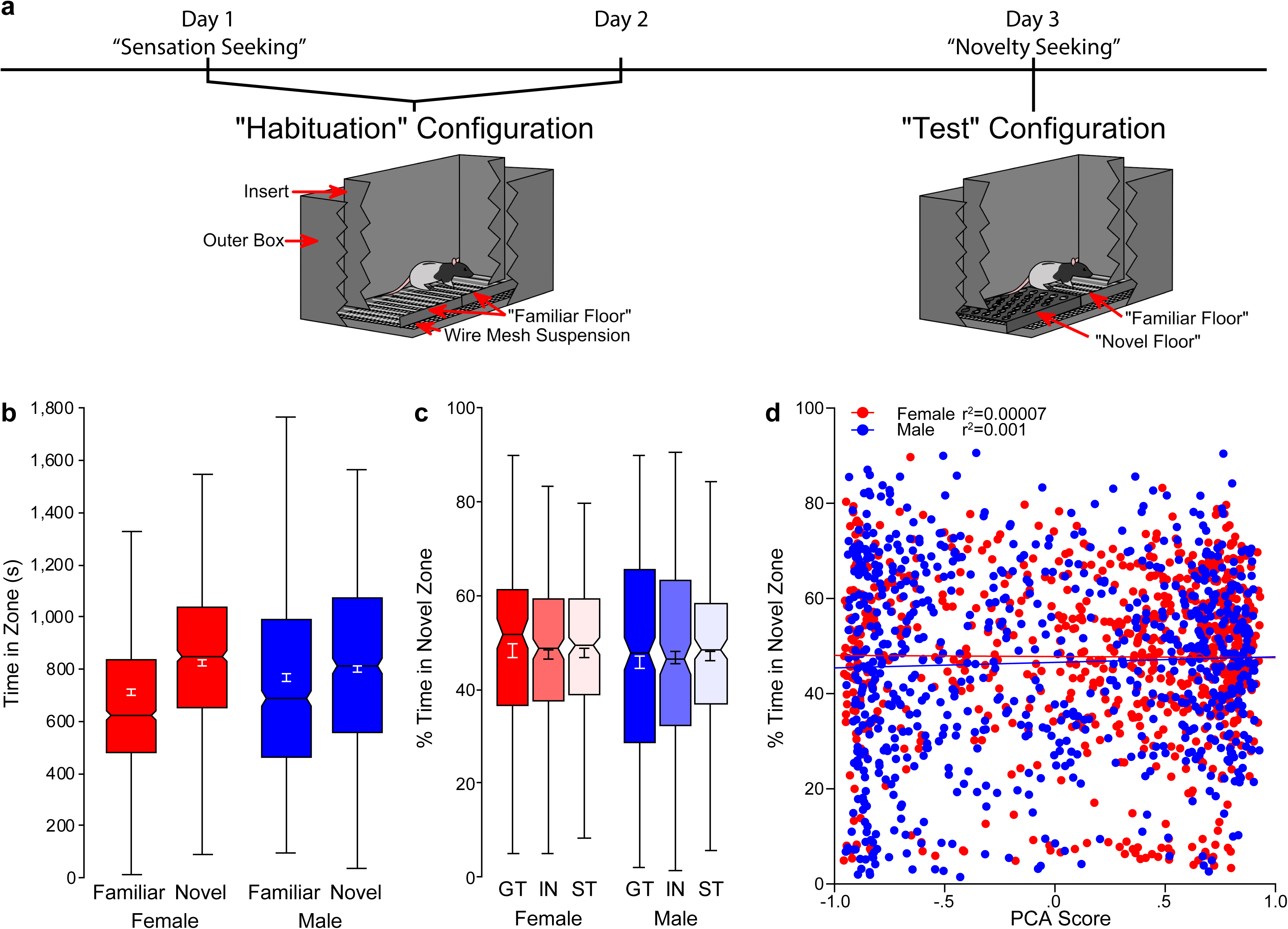
Novelty-seeking behavior. (a) Novelty-seeking paradigm timeline with schematics of the apparatus on habituation and test days. (b) Notched box plot of time spent (in seconds) in the familiar and novel zone for females (n=761) and males (n=760). (c) Notched box plot of the percent of time spent in the novel zone for females (GT (n=152), IN (n=278), ST (n=330)) and males (GT (n=287), IN (n=221), ST (n=250)). (d) Data for each individual rat’s percent time spent in the novel zone as a function of terminal PCA Index with linear regression line.

The relationship between PCA Index score and novelty-seeking behavior was also examined in subsets of the population identified by the percent of time spent in the novel zone. Those who spent <50% of the time in the novel zone were characterized as “low novelty-seekers”, and those who spent >50% of the time in the novel zone were characterized as “high novelty-seekers”. Although there was a significant (P<0.001) correlation between PCA Index score and novelty-seeking behavior in both of these populations, the r^2^ value was too low to be considered meaningful (low novelty-seekers, r^2^=0.036, high novelty-seekers, r^2^=0.043). Thus, even in the extremes of this population of heterogeneous stock rats, novelty preference and the propensity to attribute incentive salience to a food cue appear to be unrelated traits.

Given that prior reports investigating the relationship between sensation-seeking and novelty-seeking behavior are inconsistent^27-29^, we took advantage of our large sample size with both sexes represented to further examine this relationship. Similar to prior reports that used a relatively large sample size^29^, we did not find a significant correlation between these two traits when sexes were collapsed (both sexes r^2^=0.0001, p=0.694), nor when the sexes were analyzed separately (female r^2^=0.0007, p=0.456; male r^2^=0.00001, p=0.921). Furthermore, there was no relationship between these traits when examined in the subpopulations of low novelty-seekers (r^2^=0.016, p<0.001) or high-novelty-seekers (r^2^=0.035, p<0.001); nor in low-responders (r^2^=0.0004, p=0.579) or high-responders (r^2^=0.001, p=0.472).

### Principal components analysis

To determine whether the traits described above could be reduced to fewer dimensions that might better capture the variance in behavioral outcome measures, principal components analysis was performed. When the entire population was included in this analysis, the behavioral variables were reduced to two factors that, together, account for ∼63% of the variance in behavior (Figure 7 and Supplemental Table 1). Factor 1, which accounts for ∼38% of the overall variance, has strong (>0.7) loadings from PCA Index Score (0.80) and Incentive Value Index (0.82), with a weaker (0.44) and perhaps non-significant loading from sensation-seeking behavior. In contrast, Factor 2, which is orthogonal to Factor 1, accounts for 25% of the variance, and is comprised largely of a single variable: novelty-seeking behavior (loading = 0.98). Taken together, these data are largely in agreement with the regression analyses reported above, demonstrating a strong relationship between two indices reflective of the incentive motivational value of a reward cue, with novelty-seeking behavior representing an entirely separate dimension of behavioral variability. When principal components analysis was conducted separately for each sex and phenotype, the same pattern of factor loadings is apparent for both males and females and for ST and IN responders (see Supplemental Table 1). For GT, however, a slightly different picture emerges, such that Incentive Value Index (0.77) and sensation-seeking (0.75) load strongly onto Factor 1; whereas Factor 2 captures the PCA Index Score (0.71) and novelty-seeking (0.73) behavior. The relationship between the variables also changes within phenotypes when males and females are considered separately (see Supplemental Table 1). While the principal components analysis does not reveal much more than the regression analyses described above, it does highlight the fact that the relationship between traits can change in a phenotype- and sex-dependent manner.

**Figure 7.**
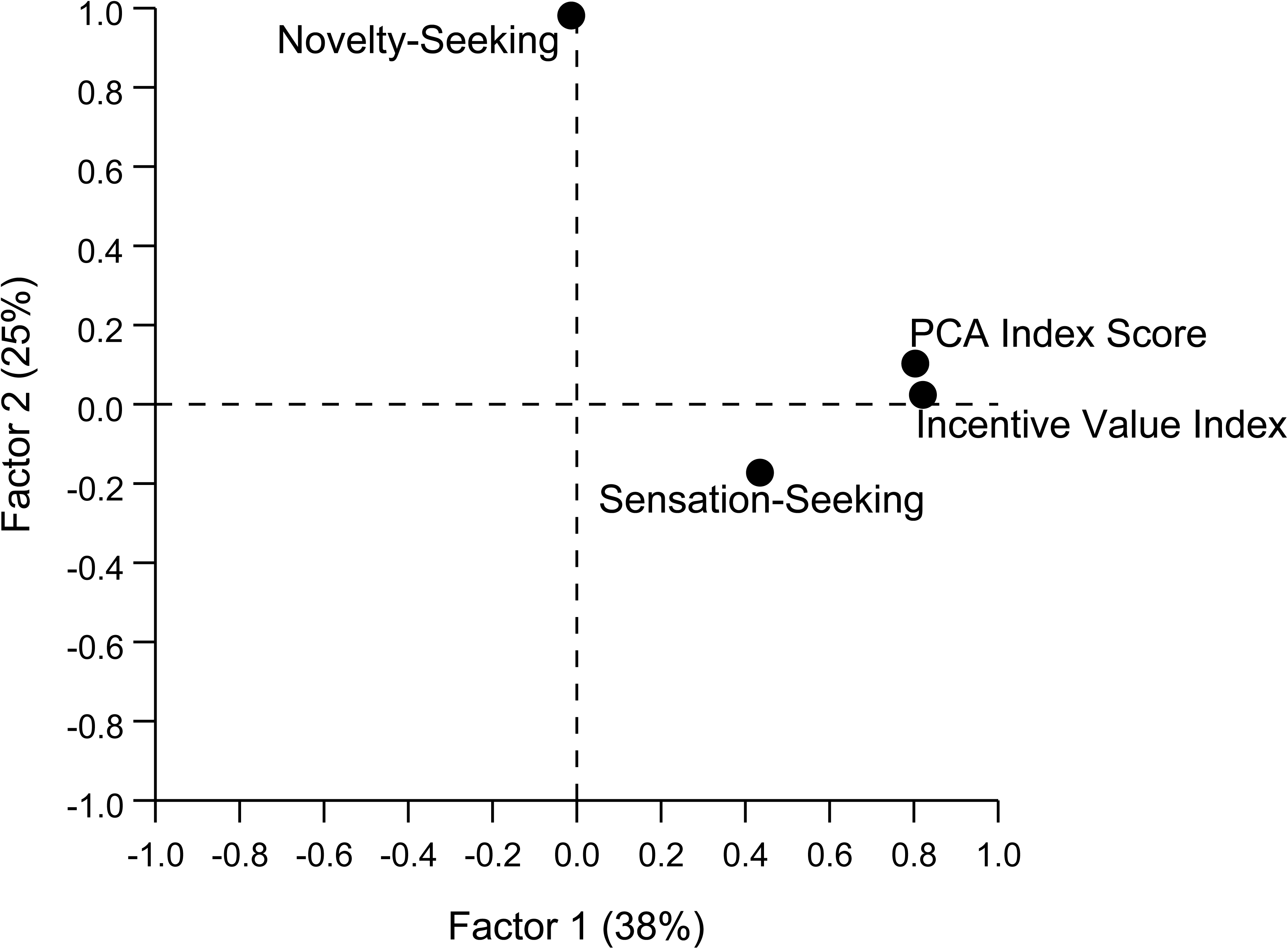
Principal components in rotated space. Principal components analysis of the relationship between: PCA Index Score, Incentive Value Index, Sensation Seeking, and Novelty Seeking. The two extracted factors cumulatively explained ∼63% of the variance in behavior. PCA Index Score and Incentive Value Index loaded strongly (>0.7) onto Factor 1, which accounts for 38% of the total variance. Additionally, Sensation Seeking loaded onto this factor to a lesser degree (0.44). Factor 2, which accounts for 25% of the variance, is comprised largely of Novelty Seeking (0.98).

## Discussion

The primary aim of the present study was to determine the relationship between individual differences in the propensity to attribute incentive salience to a reward-cue, as assessed by sign-tracking behavior, and two other traits that have been related to susceptibility to addiction, sensation-seeking and novelty-seeking behaviors. We exploited a large sample (N=1,598) of a uniquely heterogeneous strain of rats^33^ to examine the relationship between these traits. There were no meaningful correlations between the propensity to attribute incentive salience to a reward cue, sensation-seeking nor novelty-seeking behavior for either male or female rats. There were, however, novel correlative relationships identified for multiple measures of incentive salience attribution and sex differences revealed for a number of the outcome measures.

The ability of a reward-paired cue to elicit approach behavior is one of the fundamental properties of an incentive stimulus^5^; that is, a cue that has been transformed into a “motivational magnet”^5,37^ as a function of incentive salience attribution. Based on this notion, our earlier work characterized rats as sign- or goal-trackers based strictly on the number of contacts with the lever-cue upon its presentation^9,11^, but more recently we have used the PCA Index to identify sign- and goal-trackers^35^. Rather than relying on a single measure, the PCA Index incorporates the number, latency and probability to contact the lever-cue vs. the food cup^35^. Here, we show, for the first time, that this metric differs significantly between the sexes. In females, the PCA Index score is biased towards sign-tracking, both over training and as reflected in the terminal PCA Index score, relative to males (Figure 2a, b, c). In a prior study^36^ with Sprague-Dawley rats, we reported only modest sex differences, such that female sign-trackers tended to acquire their conditioned response more rapidly than males of the same phenotype. In the current study, however, sex effects were most pronounced during acquisition in intermediate responders on measures of sign-tracking behavior, and in goal-trackers on measures of goal-tracking behavior. In both cases, females showed enhanced responding relative to males. These discrepant findings are not surprising given the smaller sample sizes in our prior work (i.e. n=8-16 per sex per phenotype) and different rat strains used^36^. However, it is not clear whether the different findings are because the sex differences are dependent on genetic background, or simply a function of a small sample size in earlier studies. We suspect the latter, as there is abundant literature suggesting that the “typical” sample size used in behavioral neuroscience research will often result in non-reproducible results^38^. Furthermore, it should be noted that a similar trend was observed even in the prior study, and that intermediate responders were not included in that analysis^36^. With respect to the current findings, we speculate that the apparent sex differences may be due, at least in part, by greater activity levels in females relative to males, because females also exhibited greater responding at the food cup during the intertrial intervals, relative to males, although it is unlikely this fully accounts for the sex difference. Ongoing studies are investigating the biological bases of these effects.

A second fundamental property of an incentive stimulus is that it itself becomes an object of desire, in that an individual will work for the stimulus alone^4,5,9^. This property of an incentive stimulus is typically assessed using a conditioned reinforcement test, in which it is determined whether a rat will learn a new instrumental response for presentation of the conditioned stimulus alone, which, in this case is the lever-cue. Importantly, during the conditioned reinforcement test, food reward is absent and the reinforcer is the lever. Although all rats responded more into the port that resulted in presentation of the lever, sign-trackers did so to a greater extent than intermediate responders or goal-trackers (Figure 4b), which is consistent with our prior reports^9,35^. Also similar to our previous report^36^, we found that females showed greater responding for presentation of the lever relative to males (Figure 4a), and this is true even when greater responding at the “inactive” port is accounted for (Figure 4b). During the conditioned reinforcement test we also measured responses directed towards the lever upon its brief presentation following an instrumental response. Consistent with previous reports^9^, sign-trackers interacted with the lever more than intermediate responders or goal-trackers. This held true for both sexes, but females had a tendency to make more lever deflections than males (Figure 4c). Taken together, these data support the notion that the lever acts as a more effective conditioned reinforcer for sign-trackers, and suggest that the secondary reinforcing properties of the lever may be enhanced for females compared to males.

We previously showed that terminal PCA Index score is an effective predictor of the conditioned reinforcing properties of the lever-cue^35^, which is to be expected as both reflect the incentive motivational value of the conditioned stimulus. The current analyses extend these findings, demonstrating that the terminal PCA Index score is a much stronger predictor of the number of lever deflections during the conditioned reinforcement test than the number of instrumental responses in the nose port (i.e. A-I^35^). These findings underscore the need to incorporate the number of deflections when considering the conditioned reinforcing properties of the lever, as relying solely on nose port responding underestimates the incentive value. That is, the enhanced interest in the lever exhibited by STs competes with responding in the nose port, because they are drawn towards the lever. Thus, the number of nosepokes underestimates the incentive value of the lever to a greater degree in STs than GTs. To account for this, we generated a novel metric, the “Incentive Value Index”, which is calculated from responses into both nose ports and lever deflections during the conditioned reinforcement test. As expected, the Incentive Value Index is greater in STs compared to GTs and intermediate responders; and females have a higher Incentive Value Index compared to males. The Incentive Value Index is correlated with the terminal PCA Index for both sexes, accounting for about 15% of the variance in females and about 19% of the variance in males. Importantly, when the ability of cue presentation to promote instrumental responding is assessed using a different paradigm, wherein the competition between instrumental responding and approach to the lever-CS is not present, the relationship between PCA Index and cue-evoked responding is greater, accounting for 25% of the variance in behavior^39^. This further supports the notion that we may be underestimating the relationship between the two indices of incentive value as a function of the experimental paradigm. Taken together, these data capture the ability of the CS to act as a more effective “motivational magnet” in sign-trackers and in females, and highlight the need to assess interaction with the CS as a critical component of the conditioned reinforcement test.

The propensity to attribute incentive salience to reward cues has previously been associated with individual differences in impulsive behavior^31,40^, responsivity to aversive stimuli^41^, attentional control^42^, and susceptibility to cue- and drug-induced reinstatement of drug-seeking behavior following limited drug exposure and abstinence^14,15^. Here we examined the relationship between this trait and two others that have been associated with addiction-related behaviors. Locomotor response to an inescapable novel environment or “sensation-seeking” behavior was first described as a trait relevant to addiction liability in rodents by Piazza and colleagues who, in 1989, showed that individual differences in activity levels in a novel environment could predict the initial tendency to take drugs^23^. That is, those that showed the highest activity, or high-responders (HR), acquired drug self-administration at a faster rate relative to those that exhibited lower levels of activity, or low-responders (LR). In contrast to other reports^27^, we have previously shown that locomotor response to novelty is not correlated with the propensity to attribute incentive salience to reward cues in a population of outbred Sprague-Dawley rats^1^, and the current findings, are in agreement. That is, using a large sample of heterogeneous stock rats, we found that the correlation between “sensation-seeking” and PCA Index was too small to be considered meaningful, accounting for less 0.1% of the variance in either sex. In addition, there was not a significant difference in “sensation-seeking” behavior between phenotypes. The lack of a relationship between “sensation-seeking” behavior and incentive salience attribution in this large population of outbred animals is intriguing, given that these traits seem to have been co-selected in rats that are bred for extreme differences in locomotor response to a novel environment^31^. That is, selectively bred high-responder (bHR) rats are almost always sign-trackers; whereas selectively bred low-responder (bLR) rats are almost always goal-trackers. Yet, in the current dataset, even when only the extremes of the population were assessed, we did not observe a significant relationship between these traits. Importantly, however, the bHR rats, exhibit a unique pattern of addiction-related traits that do not appear to be related in outbred animals**^26,31,32,43^**. This is, perhaps, not too surprising, as traits that are genetically unrelated could diverge by chance between a high- and low-selected line^44^.

In addition to sensation-seeking behavior, we assessed novelty-seeking behavior or novelty place preference, as this too has been associated with addiction liability^26^. Novelty-seeking behavior was previously reported to be positively correlated with sign-tracking behavior^27^, but we did not observe this relationship in the current study. Although there was a preference for the novel zone of the testing chamber relative to the familiar, the degree of preference was comparable between phenotypes and sexes, and there was not a significant correlation between terminal PCA Index score and zone preference. It should be noted that our experimental design for this test was somewhat unconventional, and quite different from that previously used to assess the relationship between novelty-seeking behavior and the propensity to attribute incentive salience to reward cues^27^. Beckmann et al., previously used a three-compartment chamber that consisted of a novel, familiar and neutral zone that was physically divided and differentiated by both floor type and color^27^. In contrast, the apparatus used in the current study consisted of a single chamber with two distinct floor types. The “familiar” zone always consisted of a grid floor, and the “novel” zone consisted of a floor with holes in it. It was not a counterbalanced design because the grid floor was comparable to that used during Pavlovian training and we wanted to ensure that the novel zone was indeed novel. Importantly, in a prior study using the same chambers^45^, there was no bias for one floor type relative to the other in rats that were previously exposed to Pavlovian training (Paul Meyer, personal communication), as they were in the current study. Thus, we believe that novelty-seeking behavior was adequately captured with the current design. It is also important to note, however, that the dependent variables used to reflect the propensity to sign-track differed between the current study and that reported by Beckmann et al. While we relied on the Pavlovian Conditioned Approach Index to capture the tendency to sign-track, Beckmann et al. used the “percent of trials with a sign-tracking response” ^27^. One could argue that reliance on a single variable to reflect such a complex trait may lead to erroneous conclusions and the inability to replicate across batches of animals^35^. Thus, while it is likely that differences in the testing procedures, outcome measures and sample size^38^ might have contributed to the discrepant findings between the current report and those of Beckmann et al., we are confident that in this large sample of heterogeneous stock rats novelty-seeking and the propensity to attribute incentive salience to reward cues are two distinct traits. This notion is further supported by the principal components analysis, as shown in Figure 7. Furthermore, as novelty-seeking behavior has specifically been associated with the transition to compulsive drug use^26^, and the propensity to attribute incentive salience to reward cues with the tendency to relapse^15^, these data highlight distinct paths of addiction vulnerability, with dissociable traits contributing to the different phases of addiction.

The current study represents the first to investigate the relationship between addiction susceptibility traits using such a large sample size. We were fortunate to exploit a large sample of a uniquely heterogeneous rat population that is concurrently being used for other investigations that require such sample sizes^46^. The reported findings uncover novel relationships between multiple measures of incentive salience attribution, led to a new metric that captures “incentive value”, and revealed sex differences that were not previously apparent. Moreover, this work highlights the importance of sample size and effect size when interpreting results, as relationships that were previously reported to exist between traits using small sample sizes, were non-existent in our sample of ∼1,600 rats. While we fully recognize and appreciate the obstacles that preclude the utilization of such large sample sizes in behavioral neuroscience research, the results underscore the need for caution when interpreting relationships identified with relatively small samples^38,47^.

## Methods and Materials

### Subjects

Subjects were 799 male and 799 female Heterogeneous Stock (N: NIH-HS) rats provided by a breeding colony maintained at the Medical College of Wisconsin (Dr. Leah Solberg Woods, now at Wake Forest School of Medicine). The HS strain was established at the National Institute of Health (NIH) using eight inbred founder strains that were genetically and phenotypically diverse^48^. Genetic heterogeneity has been maintained by the Solberg Woods’ lab using a random breeding scheme that takes into account the kinship coefficient between animals, which minimizes inbreeding and maximizes recombination of genetic loci across each litter^33^. The colony has been maintained in this way using 64 breeder pairs since 2013.

Rats arrived at the University of Michigan at approximately 35 days of age. They were triple housed with members of the same sex on a 12-hour reverse light cycle (lights off at 0730 h). Food and water were freely available in the home cage for the duration of the experiment; that is, the animals were never food deprived. Behavioral testing began at approximately 60 days of age, and all experimentation was conducted during the dark phase between the hours of 0900 h and 1600 h. All experiments followed the principals of laboratory animal care specified by “Guidelines for the Care and Use of Mammals in Neuroscience and Behavioral Research” National Research Council (2003) and all procedures were approved by the Institutional Animal Care and Use Committee at the University of Michigan.

### Pavlovian conditioned approach (PCA)

#### Apparatus

Pavlovian conditioning occurred in standard Med Associates (St. Albans, VT) test chambers (30.5 x 24.1 x 21 cm) which were located in sound-attenuating cabinets with a ventilating fan to mask background noise. Each chamber contained a food cup, located 3 cm above the stainless-steel grid floor on the center of one wall. Banana-flavored food pellets (45 mg, BioServe, #F0059, Frenchtown, NJ, USA) were delivered into the food cup via an automatic pellet dispenser. Head entries were detected by breaks in an infrared photobeam located inside the food cup. A retractable, backlit, metal lever was placed either to the left or the right of the food cup (counterbalanced) and located 6 cm above the grid floor. Lever deflections were recorded when the lever was deflected with a minimum 10-g force. A red house light was located on the top of the wall opposite the food cup and lever, and illuminated for the duration of the session. All data were collected using MED-PC IV software, and extracted using Med-PC to Excel.

#### Pre-training

Rats were given roughly 20 banana-flavored food pellets in their home cage for two days immediately prior to pre-training in order to familiarize the rats with the food reward to be used during training. Pre-training occurred in the same Med Associates chambers where they would subsequently undergo Pavlovian training. The pre-training consisted of an approximately 12.5-minute period during which 25 banana-flavored pellets were delivered on a variable time (VT) 30-second schedule (time varied between 0 and 60 seconds). During this session food cup entries were recorded and it was confirmed that rats were reliably retrieving all of the food pellets.

#### Pavlovian conditioned approach training

After pre-training, rats underwent Pavlovian conditioned approach (PCA) training. One session was conducted daily for 5 days. Each PCA session consisted of 25 trials beginning with the presentation of an illuminated lever (which served as the conditioned stimulus, CS) for 8 seconds, immediately followed by the delivery of a banana-flavored food pellet (unconditioned stimulus, US) into the adjacent food cup. Each CS-US pairing occurred within a VT 90-second schedule (time varied between 30 and 150 seconds). The number of lever deflections, head entries into the food cup during CS presentation, and head entries into the food cup during the inter-trial intervals were recorded.

For each session, the total number of lever and food cup entries, the average latency from lever extension to lever deflection or food cup entry (in seconds, maximum 8), and probability of lever deflection and food cup entry during each trial was calculated. These metrics were combined into a PCA Index score composed of: response bias [(total lever contacts – total food cup contacts) ÷ (sum of total contacts)], probability difference score [Prob_(lever)_ – Prob_(food cup)_], and latency difference score [-(lever contact latency – food cup entry latency) ÷ 8]. These three measures were then averaged together to create the PCA Index score, which ranges from -1 to +1, with -1 being an exclusive preference for the food cup and +1 being an exclusive preference for the lever^35^. The PCA index scores for the final two sessions of training (4 and 5) were averaged into a terminal score which provided a single measure of Pavlovian conditioned approach for each rat. Based on their terminal PCA score, each individual rat was assigned a behavioral phenotype. Rats with a score below -0.5 were classed as goal-trackers (GTs) and rats with scores above 0.5 were classed as sign-trackers (STs). Rats with scores between -0.5 and 0.5 were considered intermediate responders (IN). These phenotype groups were further subdivided by sex for statistical analyses.

### Conditioned reinforcement

One day following the final Pavlovian conditioning session, rats were exposed to a conditioned reinforcement test to evaluate the reinforcing properties of the lever-CS. Conditioned reinforcement occurred in the same Med Associates test chambers described above; however, the chambers were rearranged such that the food cup was removed and the lever was moved to the center of the wall in its place. Two nose poke ports, equipped with infrared head entry detectors, were placed on the wall, to the left and right of the lever. Nose pokes into the “active” port, located where the lever had been previously, resulted in a 2-second presentation of the lever. Nose pokes into the “inactive” port had no programmed consequences. The conditioned reinforcement test lasted 40 minutes. The number of pokes into the active and inactive ports, and the number of lever deflections were recorded. The difference between active and inactive nose pokes (A-I) was also derived from these data. This metric was used for correlational analyses, and allowed us to account for potential differences in activity at the inactive nose port that were, presumably, independent of the reinforcing properties of the lever. In addition, we also analyzed the “Incentive Value Index”, which was calculated using the following formula: ((responses in active port – responses in inactive port) + lever deflections)). As described above, the Incentive Value Index was devised as a means to account for the fact that relying solely on the number of operant responses during a test for conditioned reinforcement likely underestimates the incentive value of the lever-CS.

### Sensation-seeking and novelty-seeking

#### Apparatus

The tests for sensation-seeking and novelty-seeking took place in 12 chambers made from expanded PVC and comprised of an outer box (68.58 x 33.02 x 66.04 cm) and a smaller insert box (45.72 x 15.24 x 55.88 cm) (see Figure 6a). Each chamber had a wire mesh suspended across the bottom of the outer box, upon which interchangeable floors could be placed. The insert box was arranged on top of these floors, creating an inescapable chamber. A camera (CVC-130R, Speco Technologies, Amityville, NY, USA) was suspended approximately 18 cm above the center of each insert to record locomotor activity and videos were analyzed using Noldus (Leesburg, VA, USA) Ethovision motion-tracking software.

Two days following the conditioned reinforcement test, rats were placed into the testing apparatus. As described below and in Figure 6, the first exposure allowed us to assess “sensation-seeking”; while the last exposure was the “novelty-seeking” test.

#### Sensation-seeking (habituation)

The first 30-min exposure to the test chamber served two purposes. First, it allowed assessment of the locomotor response to a novel environment or sensation-seeking behavior, which was measured as total distance travelled (m). Second, this first exposure served as a habituation session to what would become the “familiar” floor. The “familiar” floor was made of parallel stainless steel bars spaced 1.27 cm apart and arranged perpendicular to the long axis of the smaller insert. Rats were exposed to this grid floor for two 30-min sessions on consecutive days (see Figure 6a). During both sessions behavior was captured by the overhead cameras and locomotor activity was quantified using Ethovision.

#### Novelty-seeking test

On the third day, following the two habituation sessions, rats underwent a test session in which half of the floor was replaced with a “novel” floor composed of a solid metal plate with 1.27 cm diameter holes distributed evenly across the surface (see Figure 6a). Given the setup of the testing room and other considerations for uniformity, 1/3 of the test chambers had the “novel” floor in the opposite configuration relative to the other chambers. The composition of the “novel” hole floor was not counterbalanced, as the “familiar” (grid) floor was similar to the grid floor that the rats were previously exposed to during Pavlovian conditioned approach training. Therefore, given the rats were already familiar with a grid floor, we could not make it novel. Furthermore, preference for the grid vs. hole floor was assessed in a prior study^45^ using the same apparatus following Pavlovian conditioned approach training, and there were no significant differences (t(30) = 0.89, P=0.38) in the amount of time spent on either floor type (Paul Meyer, personal communication).

On the test day, rats were again placed into the chambers for 30 minutes (1800 sec) and their locomotion was captured by the cameras suspended overhead. Videos were analyzed using Ethovision and the time spent in each zone (“Familiar” and “Novel”) was recorded. When a rat was neither fully in (i.e. with all 4 paws) the “familiar” or “novel” zone, it was considered to be in the “neutral” zone. Thus, for a given rat, the total time spent in either the “familiar” or “novel” zone will not always add up to 1800 sec, due to time spent in the “neutral” zone. A “novelty place preference score” was calculated as the percentage of the total session time each rat spent in the novel zone, and this metric was used for correlational analyses.

### Statistics

The primary purpose of the present experiments was to determine whether or not there are significant relationships between an individual’s propensity to attribute incentive salience to a reward cue, assessed by Pavlovian conditioned approach behavior, and the expression of sensation-seeking or novelty-seeking behavior. While 1,598 (799 males, 799 females) rats completed PCA training and were phenotyped according to their PCA Index score, the number of valid observations on subsequent measures was often less, as some data points were lost due to equipment malfunction or an ill animal. If subjects made no responses during a testing period, but it was determined that the apparatus functioned properly and the animal was in good health during the test, the measurement was recorded as zero and the subject was included in the analyses.

All ANOVA,regression analyses and principal components analyses were conducted using SPSS 24. The effect of phenotype (i.e. ST, GT, IN) and sex were assessed for each metric of incentive salience attribution, sensation-seeking and novelty-seeking behavior using two-way ANOVAs, or with linear mixed models when session was included as a covariate. When a significant effect (p<0.05) was revealed, main effects and interactions were further analyzed using Bonferroni corrections. Given the large sample size and potentially inflated statistical power, we also report effect sizes for pairwise comparisons using Cohen’s d^49^. Cohen suggested^50^ that effect sizes less than 0.2 be considered “small”, and those greater than 0.8 considered “large”. In the current dataset, however, we take a relatively conservative approach and consider the nature of each measurement and the supporting test statistics to determine whether a given effect size constitutes a meaningful result.

The predictive relationship between the propensity to attribute incentive salience to reward cues, sensation-seeking and novelty-seeking behavior was assessed using linear regression. For all regressions, we also assessed the effect of sex using a one-way ANOVA, with the predicting variable (PCA Index) as a covariate. This allowed us to assess interactions between sex and PCA Index to determine if the predictive value of PCA Index on other metrics differed between males and females.

Principal components analysis was used to determine whether the traits of interest could be reduced to fewer dimensions and to identify underlying constructs. The behavioral variables included in this analysis were the same as those included in regression analysis as described above. That is, 1) Pavlovian Conditioned Approach Index, 2) Incentive Value Index, 3) Sensation-seeking behavior (i.e. distance travelled) and 4) Novelty-seeking behavior (i.e. % time in novel zone). The number of factors was determined using a minimum Eigenvalue criterion of 1 and resulting principal components were rotated using the Varimax method.

### Image processing

Notched box plots were created in Microsoft Excel using the XLSTAT Free add-on (Descriptive Statistics grouped box plot with notched option). Line plots, histograms, and scatterplots were made using SPSS syntax and edited (axes range standardized, colors and fit lines applied to scatterplots) in SPSS chart editor. Adobe Illustrator was used to compile each figure and to construct the schematics for the novelty-seeking procedure (Figure 6) and the overall experimental timeline (Supplemental Figure 1). The illustrations for the novelty-seeking apparatus were made in Moho 12.

Final processing of each figure was conducted in Adobe Illustrator. The specific processing manipulations are as follows: Font style/size and position of axes labels and numbers were standardized, axes label text was edited to make all figures consistent, colors were added/edited on each figure, notches on the notched box plots were deepened (median line was shortened horizontally) to improve clarity, standard errors were layered below line plot markers, charts were resized (maintaining aspect ratio) to be uniform for each figure, individual boxes on the notched box plots were moved horizontally to compress the size of each chart and clarify groups, legends were constructed where appropriate, r^2^ values were indicated for each scatterplot.

### Data availability

The datasets generated and/or analyzed during the current study are available from the corresponding author on reasonable request.

## Acknowledgements

Funding for this work was provided by a Center of Excellence grant (P50DA0377844) from the National Institute on Drug Abuse branch of the National Institutes of Health, for which Dr. Abraham Palmer is the Principal Investigator and Drs. Shelly Flagel and Terry Robinson Co-Principal Investigators on Project 1. We would also like to acknowledge Dr. Joshua Haight for his dissertation work in the Flagel Lab, which led us to evaluate the Incentive Value Index in the current study.

## Contributions

A.R.H. and A.P.H. conducted the behavioral assays, compiled and analyzed the datasets, constructed figures and tables, and drafted the Introduction, Methods, and Results sections of the manuscript. L.C.S.W and K.H. maintained the HS rat colony and provided the rats for these experiments. A.A.P. generated some of the ideas for the project and analyses and served as the Principal Investigator for the NIDA Center of Excellence (P50 DA0377844) grant which funded the work. S.B.F. conceived the idea for the project, designed and oversaw the experiments, wrote the Discussion section, and edited and revised other portions of the manuscript. T.E.R. conceived the idea for the project and revised and edited the manuscript. All authors read and approved the final manuscript.

---

**Supplemental Figure 1.**
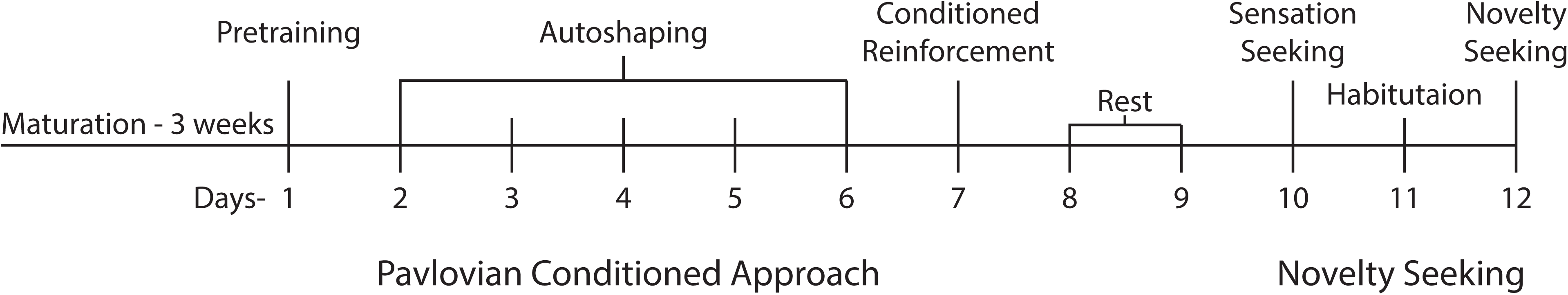
Experimental timeline. Timeline of experiment from rats’ arrival to the end of the novelty-seeking test. Rats arrived ∼35 days of age and were allowed ∼3 weeks to acclimate and mature to adulthood. Behavioral testing commenced when they were ∼60 days old and was completed by the time they were ∼75 days old.

**Supplemental Table 1.**
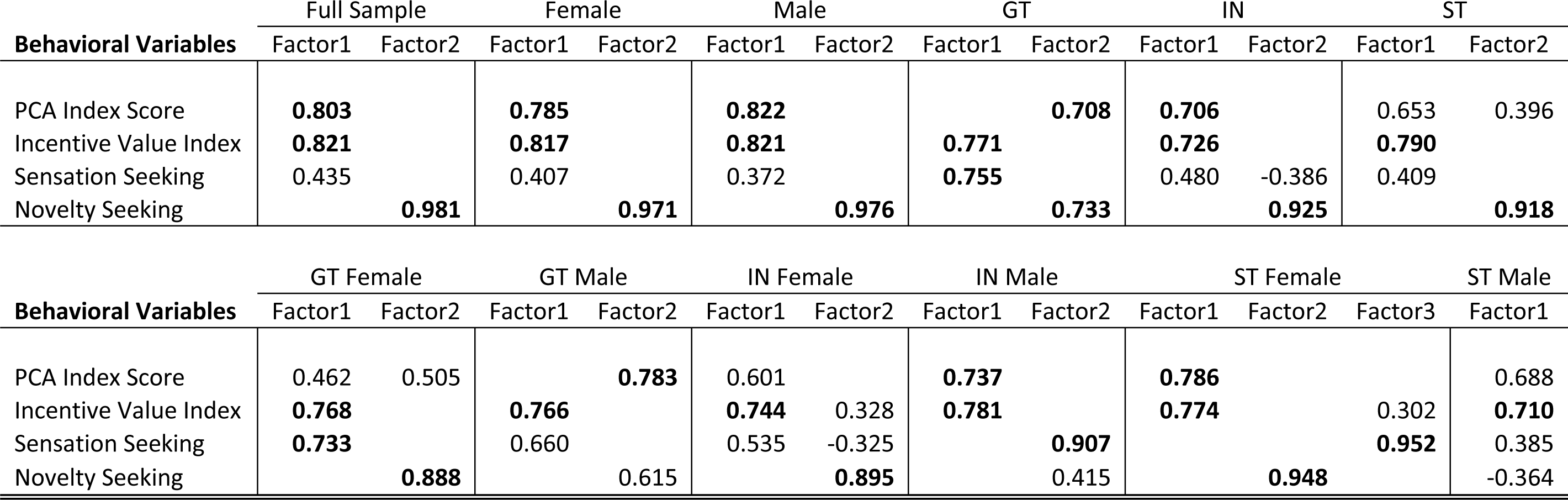
Principal components analysis. Factor loadings from the rotated component matrix for the entire population (i.e. full sample) and for each sex and phenotype considered separately. Loadings >0.7 are shown in bold. Those for the full sample correspond to Figure 7.

## References

1 Flagel, S. B., Akil, H. & Robinson, T. E. Individual differences in the attribution of incentive salience to reward-related cues: Implications for addiction. Neuropharmacology 56, 139–148, doi:10.1016/j.neuropharm.2008.06.027 (2009).

2 Ehrman, R. N., Robbins, S. J., Childress, A. R. & Obrien, C. P. Conditioned-responses to cocaine-related stimuli in cocaine abuse patients. Psychopharmacology 107, 523–529, doi:Doi 10.1007/Bf02245266 (1992).

3 Obrien, C. P., Childress, A. R., Mclellan, A. T. & Ehrman, R. Classical-conditioning in drug-dependent humans. Ann Ny Acad Sci 654, 400–415, doi:DOI 10.1111/j.1749-6632.1992.tb25984.x (1992).

4 Berridge, K. C. Reward learning: Reinforcement, incentives, and expectations. Psychology of learning and motivation: Advances in research and theory 40, 223–278 (2001).

5 Cardinal, R. N., Parkinson, J. A., Hall, J. & Everitt, B. J. Emotion and motivation: the role of the amygdala, ventral striatum, and prefrontal cortex. Neurosci Biobehav R 26, 321–352, doi:Pii S0149-7634(02)00007-6 Doi 10.1016/S0149-7634(02)00007-6 (2002).

6 Stewart, J., Dewit, H. & Eikelboom, R. Role of unconditioned and conditioned drug effects in the self-administration of opiates and stimulants. Psychol Rev 91, 251–268, doi:Doi 10.1037/0033-295x.91.2.251 (1984).

7 Di Ciano, P. & Everitt, B. J. Conditioned reinforcing properties of stimuli paired with self-administered cocaine, heroin or sucrose: implications for the persistence of addictive behaviour. Neuropharmacology 47, 202–213, doi:10.1016/j.neuropharm.2004.06.005 (2004).

8 Robinson, T. E. & Berridge, K. C. The neural basis of drug craving - an incentive-sensitization theory of addiction. Brain Res Rev 18, 247–291, doi:Doi 10.1016/0165-0173(93)90013-P (1993).

9 Robinson, T. E. & Flagel, S. B. Dissociating the predictive and incentive motivational properties of reward-related cues through the study of individual differences. Biol Psychiat 65, 869–873, doi:10.1016/j.biopsych.2008.09.006 (2009).

10 Boakes, R. A. Performance on learning to associate a stimulus with positive reinforcement. Operant-Pavlovian Interactions (ed. Davis, H., Hurwitz, H.), 67-97 (1977).

11 Flagel, S. B., Watson, S. J., Robinson, T. E. & Akil, H. Individual differences in the propensity to approach signals vs goals promote different adaptations in the dopamine system of rats. Psychopharmacology 191, 599–607, doi:10.1007/s00213-006-0535-8 (2007).

12 Zener, K. The significance of behavior accompanying conditioned salivary secretion for theories of the conditioned response. Am J Psychol 50, 384–403, doi:Doi 10.2307/1416644 (1937).

13 Hearst, E., Jenkins, H. Sign-tracking: The stimulus-reinforcer relation and directed action. (1974).

14 Saunders, B. T. & Robinson, T. E. Individual variation in the motivational properties of cocaine. Neuropsychopharmacol 36, 1668–1676, doi:10.1038/npp.2011.48 (2011).

15 Saunders, B. T., & Robinson, T. E. A cocaine cue acts as an incentive stimulus in some but not others: implications for addiction. Biol Psychiat 67, 730–736 (2010).

16 Ayduk, O. et al. Regulating the interpersonal self: Strategic self-regulation for coping with rejection sensitivity. J Pers Soc Psychol 79, 776–792, doi:Doi 10.1037/0022-3514.79.5.776 (2000).

17 Franques, P. et al. Sensation seeking as a common factor in opioid dependent subjects and high risk sport practicing subjects. A cross sectional study. Drug Alcohol Depen 69, 121–126, doi:Pii S0376-8716(02)00309-5 Doi 10.1016/S0376-8716(02)00309-5 (2003).

18 Franques, P., Auriacombe, M. & Tignol, J. Addiction and personality. Encephale 26, 68–78 (2000).

19 Khan, A. A., Jacobson, K. C., Gardner, C. O., Prescott, C. A. & Kendler, K. S. Personality and comorbidity of common psychiatric disorders. Brit J Psychiat 186, 190–196, doi:DOI 10.1192/bjp.186.3.190 (2005).

20 Kreek, M. J., Nielsen, D. A., Butelman, E. R. & LaForge, K. S. Genetic influences on impulsivity, risk taking, stress responsivity and vulnerability to drug abuse and addiction. Nat Neurosci 8, 1450–1457, doi:10.1038/nn1583 (2005).

21 Masse, L. C. & Tremblay, R. E. Behavior of boys in kindergarten and the onset of substance use during adolescence. Arch Gen Psychiat 54, 62–68 (1997).

22 Zuckerman, M., & Cloninger, C. R. Relationships between Cloninger’s, Zuckerman’s, and Eysenck’s dimensions of personality. Personality and Individual Differences 21, 283–285 (1996).

23 Piazza, P. V., Deminiere, J. M., Lemoal, M. & Simon, H. Factors that predict individual vulnerability to amphetamine self-administration. Science 245, 1511–1513, doi:DOI 10.1126/science.2781295 (1989).

24 Hughes, R. Behaviour of male and female rats with free choice of two environments differing in novelty. Animal Behaviour 16, 92–96 (1968).

25 Belin, D., Mar, A. C., Dalley, J. W., Robbins, T. W. & Everitt, B. J. High impulsivity predicts the switch to compulsive cocaine-taking. Science 320, 1352–1355, doi:10.1126/science.1158136 (2008).

26 Belin, D., Berson, N., Balado, E., Piazza, P. V. & Deroche-Gamonet, V. High-Novelty-Preference Rats are Predisposed to Compulsive Cocaine Self-administration. Neuropsychopharmacol 36, 569–579, doi:10.1038/npp.2010.188 (2011).

27 Beckmann, J. S., Marusich, J. A., Gipson, C. D. & Bardo, M. T. Novelty seeking, incentive salience and acquisition of cocaine self-administration in the rat. Behav Brain Res 216, 159–165, doi:10.1016/j.bbr.2010.07.022 (2011).

28 Lukkes, J. L., Thompson, B. S., Freund, N. & Andersen, S. L. The developmental inter-relationships between activity, novelty preferences, and delay discounting in male and female rats. Dev Psychobiol 58, 231–242, doi:10.1002/dev.21368 (2016).

29 Pelloux, Y., Costentin, J., Duterte-Boucher, D. Differential effects of novelty exposure on place preference conditioning to amphetamine and its oral consumption. Psychopharmacology 171, 277–285 (2004).

30 Vanhille, N., Belin-Rauscent, A., Mar, A. C., Ducret, E. & Belin, D. High locomotor reactivity to novelty is associated with an increased propensity to choose saccharin over cocaine: New insights into the vulnerability to addiction. Neuropsychopharmacol 40, 577–589, doi:10.1038/npp.2014.204 (2015).

31 Flagel, S. B. et al. An animal model of genetic vulnerability to behavioral disinhibition and responsiveness to reward-related cues: Implications for addiction. Neuropsychopharmacol 35, 388–400, doi:10.1038/npp.2009.142 (2010).

32 Flagel, S. B., Waselus, M., Clinton, S. M., Watson, S. J. & Akil, H. Antecedents and consequences of drug abuse in rats selectively bred for high and low response to novelty. Neuropharmacology 76, 425–436, doi:10.1016/j.neuropharm.2013.04.033 (2014).

33 Solberg Woods, L. C. QTL mapping in outbred populations: successes and challenges. Physiol Genomics 46, 81–90, doi:10.1152/physiolgenomics.00127.2013 (2014).

34 Fitzpatrick, C. J. e. a. Variation in the form of Pavlovian conditioned approach behavior among outbred male Sprague-Dawley rats from different vendors and colonies: signtrackingvs. goal-tracking. Plos One 8, e75042 (2013).

35 Meyer, P. J. et al. Quantifying individual variation in the propensity to attribute incentive salience to reward cues. Plos One 7, doi:ARTN e38987 10.1371/journal.pone.0038987 (2012).

36 Pitchers, K. K. et al. Individual variation in the propensity to attribute incentive salience to a food cue: Influence of sex. Behav Brain Res 278, 462–469, doi:10.1016/j.bbr.2014.10.036 (2015).

37 Berridge, K. C. & Robinson, T. E. Parsing reward. Trends Neurosci 26, 581–581, doi:DOI 10.1016/j.tins.2003.09.001 (2003).

38 Button, K. S. et al. Power failure: why small sample size undermines the reliability of neuroscience (vol 14, pg 365-376, 2013). Nat Rev Neurosci 14, 444–444, doi:10.1038/nrn3502 (2013).

39 Saunders, B. T., Yager, L. M. & Robinson, T. E. Cue-Evoked Cocaine “Craving”: Role of Dopamine in the Accumbens Core. J Neurosci 33, 13989–14000, doi:10.1523/Jneurosci.0450-13.2013 (2013).

40 Lovic, V., Saunders, B. T., Yager, L. M. & Robinson, T. E. Rats prone to attribute incentive salience to reward cues are also prone to impulsive action. Behav Brain Res 223, 255–261, doi:10.1016/j.bbr.2011.04.006 (2011).

41 Morrow, J. D., Maren, S. & Robinson, T. E. Individual variation in the propensity to attribute incentive salience to an appetitive cue predicts the propensity to attribute motivational salience to an aversive cue. Behav Brain Res 220, 238–243, doi:10.1016/j.bbr.2011.02.013 (2011).

42 Paolone, G., Angelakos, C. C., Meyer, P. J., Robinson, T. E. & Sarter, M. Cholinergic control over attention in rats prone to attribute incentive salience to reward cues. J Neurosci 33, 8321–8335, doi:10.1523/Jneurosci.0709-13.2013 (2013).

43 Deroche-Gamonet, V., Belin, D. & Piazza, P. V. Evidence for addiction-like behavior in the rat. Science 305, 1014–1017, doi:DOI 10.1126/science.1099020 (2004).

44 Falconer, D. S. & Mackay, T. F. C. Introduction to Quantitative Genetics (4th Edition). (Pearson, 1996).

45 Meyer, P. J., Ma, S. T. & Robinson, T. E. A cocaine cue is more preferred and evokes more frequency-modulated 50-kHz ultrasonic vocalizations in rats prone to attribute incentive salience to a food cue. Psychopharmacology 219, 999–1009, doi:10.1007/s00213-011-2429-7 (2012).

46 Chitre, A. S. et al. Genome wide association study of body weight, body mass index, adiposity, and fasting glucose in 3,173 outbred rats. doi:doi.org/10.1101/422428 (2018).

47 Rousselet, G. A., Pernet, C. R. & Wilcox, R. R. Beyond differences in means: robust graphical methods to compare two groups in neuroscience. Eur J Neurosci 46, 1738–1748, doi:10.1111/ejn.13610 (2017).

48 Hansen, C. & Spuhler, K. Development of the National-Institutes-of-Health genetically heterogeneous rat stock. Alcohol Clin Exp Res 8, 477–479, doi:DOI 10.1111/j.1530-0277.1984.tb05706.x (1984).

49 Sawilowsky, S. S. New Effect Size Rules of Thumb. Journal of Modern Applied Statistical Methods 8, 597–599 (2009).

50 Cohen, J. Statistical power analysis for the behavioral-sciences. Percept Motor Skill 67, 1007–1007 (1988).

